# High Leaf Respiration Rates May Limit the Success of White Spruce Saplings growing in The *Kampfzone* at the Arctic Treeline

**DOI:** 10.1101/2021.08.10.455813

**Authors:** Kevin L. Griffin, Stephanie C. Schmiege, Sarah G. Bruner, Natalie T. Boelman, Lee A. Vierling, Jan U. H. Eitel

## Abstract

Arctic Treeline is the transition from the boreal forest to the treeless tundra and may be determined by growing season temperatures. The physiological mechanisms involved in determining the relationship between the physical and biological environment and the location of treeline are not fully understood. In Northern Alaska we studied the relationship between temperature and leaf respiration in 36 white spruce (*Picea glauca*) trees, sampling both the upper and lower canopy, to test two research hypotheses (H_0_). The first H_01_ is that canopy position will not influence leaf respiration. The associated alternative hypothesis (H_A_) is that the upper canopy leaves which are more directly coupled to the atmosphere will experience more challenging environmental conditions and thus have higher respiration rates to facilitate metabolic function. The second H_02_ is that tree size will not influence leaf respiration. The associated H_A_ is that saplings (stems that are 5-10 cm DBH (diameter at breast height)) will have higher respiration rates than trees (stems ≥ 10 cm DBH) since saplings represent the transition from seedlings growing in the more favorable aerodynamic boundary layer, to trees which are fully coupled to the atmosphere but of sufficient size to persist. Respiration did not change with canopy position, however respiration at 25°C was 42% higher in saplings compared to trees (3.43 ± 0.19 vs. 2.41 ± 0.14 μmol m^-2^ s^-1^). Furthermore, there were significant differences in the temperature response of respiration, and seedlings reached their maximum respiration rates at 59°C, more than two degrees higher than trees. Our results demonstrate that the respiratory characteristics of white spruce saplings at treeline are extreme, imposing a significant carbon cost that may contribute to their lack of perseverance beyond treeline. In the absence of thermal acclimation, the rate of leaf respiration could increase by 57% by the end of the century, posing further challenges to the ecology of this massive ecotone.

This paper is dedicated to the memory of James N. Siedow, Professor of Botany at Duke University. I am honored to have learned from, and to have been inspired by Jim. I will always be grateful for the time he spent helping me, the depth of education he gave me, and his steady mentoring as part of my thesis committee. His devotion to science and push for deeper knowledge of plant respiration set an example for all of us, particularly those who were lucky enough to study with him. Jim was also a memorable Father Christmas at the departmental holiday parties, always keeping us laughing with his particular brand of quick wit and sarcasm. - *Kevin L. Griffin*

## INTRODUCTION

Respiration is an ancient, fundamental metabolic process supplying the energy and carbon skeletons needed to support all living things. At the global scale this process represents a massive flux of carbon from the biosphere to the atmosphere (Li *et al*., 2018) and thus plays a critical role in the global carbon cycle. Consequently, respiration needs to be well described to predict rates of climate change (Atkin *et al*., 2015; Huntingford *et al*., 2017). While our understanding of biological and environmental controls of respiration rates remains incomplete, it is clear that temperature has a first order effect, setting up potential feedbacks between environmental change and respiratory CO_2_ release (O’Sullivan *et al*., 2013; Heskel *et al*., 2016; O’Sullivan *et al*., 2017). Understanding, quantifying and predicting rates of respiration across scales of biological organization is an important challenge for science and thus humanity.

At the landscape scale, respiration represents a significant carbon loss and offsets photosynthetic carbon gain, which may have a controlling effect on ecosystem productivity. For example, eddy covariance studies across European forests identified respiratory processes as the controlling flux determining net ecosystem carbon exchange (NEE) (Valentini *et al*., 2000). Furthermore, this work showed a clear latitudinal trend in NEE (increasing with decreasing latitude), despite a distinct lack of response in gross primary productivity, suggesting that respiratory fluxes have even stronger control on productivity in northern ecosystems. In addition, a synthesis of ecosystem model outputs from arctic and boreal ecosystems identifies both autotrophic and heterotrophic respiration as key “missing piece(s)” limiting modeling efforts in northern systems (Fisher *et al*., 2018). Uncertainty in autotrophic respiration alone can be as high as 25g C m^-2^ month^-1^, a flux that is, in some boreal locations, five times the size of the uncertainty in NEE (<5 g m^-2^ month^-1^, (Fisher *et al*., 2018). The accelerated rate of warming near the poles (Cohen *et al*., 2014), the massive carbon stores across northern biomes (Fisher *et al*., 2014) and the uncertainty in the future strength of northern latitudes as a strong carbon sink (Schuur *et al*., 2015) all point to an acute need for understanding the response of respiration to temperature in these systems.

Respiratory control of ecosystem form and function may be particularly acute at the forest tundra ecotone (FTE). The FTE represents the transition from the taiga biome to the treeless tundra at higher latitudes. This vast transition zone is circumpolar and thus represents the largest terrestrial ecological transition zone found on the planet (Callaghan, T. V. *et al*., 2002). The distinct ecology of the FTE is of intense interest as climate envelope models predict that northern treeline could move rapidly towards the arctic ocean (ACIA, 2005; Pearson *et al*., 2013; Zhang *et al*., 2013), completely displacing the tundra in some North American locations and significantly altering the ecology (and carbon balance) of massive areas. While our understanding of the precise control of the location of the FTE is incomplete, ample evidence points to temperature as the primary driver (Callaghan *et al*., 2002; Sveinbjornsson *et al*., 2002; Körner, 2012b *and references therein*). Clearly other environmental and biological factors can contribute to the maintenance of the northern treeline in specific locations and these include: soils, nutrition, drought, tree demography, herbivory, abrasion and exposure, and microtopography (Körner, 2012b *and references therein;* Maguire *et al*., 2019). However, temperature is of particular interest since it is increasing rapidly in northern ecosystems (Cohen *et al*., 2014), can have a large effect on the rate of autotrophic respiration (Heskel *et al*., 2016) and is the critical driver of respiration in ecosystem models which contain large uncertainty in the carbon balance of northern ecosystems (Fisher *et al*., 2018). This suggests a clear need for additional information regarding the temperature dependence of the physiological processes controlling northern ecosystem carbon balance including photosynthesis, respiration and cell division.

White spruce (*Picea glauca* (Monench) Voss) has a transcontinental range and is one of the most common tree species defining the FTE in North America (Sutton, 1969). Considered to be one of the hardiest coniferous species, white spruce has a suite of structural and functional traits that may help explain its persistence in the harsh FTE environment where tree growth can be limited by low winter temperatures, short growing seasons, extreme light environments and often limiting rooting volumes due to the occurrence of permafrost. These potentially adaptive traits include strong apical dominance (Nienstaedt & Zasada, 1990), prolific seed production beginning at an early age (Sutton, 1969), xylem morphology capable of desiccation tolerance (Pampuch *et al*., 2020), strong photoperiodicity of growth (Eitel *et al*., 2019), early rehydration and initiation of growth despite experiencing extremely cold winters (Eitel *et al*., 2020) and rapid photoprotection mechanisms (Maguire *et al*., 2020), among many others. The physiology of white spruce is less well characterized, but studies of both photosynthesis and respiration (*e*.*g*., Weger & Guy, 1991a; Man & Lieffers, 1997; McNown & Sullivan, 2013; Stinziano & Way, 2017; Benomar *et al*., 2018; Prud’homme *et al*., 2018) indicate the physiology of leaf carbon balance in white spruce is under strong environmental control that could contribute to the establishment and persistence of the FTE. Nevertheless, the response of respiration to temperature remains largely an enigma. A fuller understanding of autotrophic respiration, and its response to temperature in this important northern treeline species would provide mechanistic information that is critically needed to assess the rapidly changing ecology of northern forests, boreal forest carbon storage, and ultimately the location of the circumpolar FTE.

Our ability to model and predict the contribution of white spruce respiration to the structure and function of the FTE is likely to require more than just a robust quantification of average respiration rates at a common temperature. Clearly the temperature response must be described and typically knowledge of thermal acclimation would also be required. However, previous work on this species suggests that thermal acclimation of leaf physiology is limited (Stinziano & Way, 2017) (Benomar *et al*., 2018). Additional considerations include canopy position, as light acclimation has been shown to dramatically affect respiration rates in white spruce (Awada & Redmann, 2000) and intra-canopy gradients in light absorption may exist. Furthermore, the top of the canopy is often the most physiologically active portion of the crown and has been shown to affect both average respiration rates and the response of respiration to temperature (Griffin *et al*., 2001; Griffin *et al*., 2002; Whitehead *et al*., 2004). An additional concern is the potential effect of size. Young saplings can be faster growing (Brienen *et al*., 2020) and thus may require higher rates of respiration to support the metabolic costs of growth (Reich, 2014; Colesie *et al*., 2020). Furthermore, it has been suggested (Körner, 2012 a & b) that saplings (defined as being less than 10cm diameter at breast height (DBH)) are significantly more susceptible to the harsh climatic conditions existing at tree line, resulting in a high rate of mortality that is speculated to contribute to the geographic location of the FTE. Recognized as the “kampfzone” in the literature (Körner, 2012b), this transitional area is characterized by an abundance of seedlings and shrub-like ‘krummholz’ but a low number of saplings that is thought to be the result of an aerodynamic boundary layer. Within this warm, still boundary layer conditions are conducive to growth, permitting seedling regeneration and krummholz survival (Körner, 2012b). However, beyond the boundary layer saplings must survive the more variable and extreme environmental conditions which arise as their increased size couples them to the atmosphere. To our knowledge, the relationships between tree size (and thus strength of the coupling to the atmosphere) and physiological function have not been studied at the FTE. Finally identifying potential proxies for respiratory function such as leaf nitrogen concentration (e.g., Griffin et al., 2001) could substantially simplify ongoing efforts to model and track the ecology and location of the FTE.

Working at an established FTE experimental site in northern Alaska (Eitel *et al*., 2019), we test two research hypotheses. The first (H_01_) is that leaf respiration and its response to temperature are not influenced by canopy position, and the second (H_02_) is that leaf respiration and its response to temperature are not influenced by tree size (saplings vs. trees). The corresponding alternative hypotheses being that the leaves at the top of the canopy (within 1 m of the apical meristem) will have higher respiration rates but a more muted response to temperature than leaves at the bottom of the canopy (1.37 m from the ground) (H_A1_), and that saplings (<10 cm DBH) will have higher respiration rates but a more muted response to temperature than trees (> 10 cm DBH) (H_A2_). The logic for the first prediction stems from the discussion above and the concept of the kampfzone, since the top of a tree tends to be more metabolically active, and the top of the canopy is more strongly coupled to the atmosphere than the bottom of the canopy. Strong atmospheric coupling would suggest that leaves at the canopy top experience a greater range of leaf temperatures which would result in a less steep temperature response overall. Similarly, the logic for the second prediction again comes from the concept of the kampfzone, suggesting that saplings are more susceptible to the harsh climatic conditions existing at tree line, which we predict would result in higher metabolic costs and result in higher average respiration rates but a more muted response to temperature. In addition to testing these two research hypotheses, we explore differences in *T*_max_ (the maximum temperature at which respiration continues to increase) and the relationships between respiration and leaf nitrogen. We stress that rapid Arctic warming is likely to result in a significantly greater respiratory loss of carbon in trees and ultimately this puts the future of the arctic and boreal carbon sink in question. By quantifying thermally induced changes in respiratory carbon fluxes, we add critically needed mechanistic information and extend our understanding of ecosystem form and function at the FTE.

## MATERIALS AND METHODS

### Site Description and Leaf Material

Our experimental sites are located in northern Alaska, on the southern side of the Brooks Range along a 5.5 km long stretch of the Dalton Highway (67°59’ 40.92” N latitude, 149°45’15.84” W longitude) and are described more fully in Eitel *et al*., (2019). *Picea glauca* (white spruce) is the only tree species at these sites, with the only exceptions being the rare black spruce tree (*Picea mariana* (Mill.) Britton, Sterns & Poggenb.), or the more common smaller multi-stemmed red alder (*Alnus rubra* Bong.). The understory contains deciduous shrubs, sedges, forbs and mosses, which are also common in the treeless tundra just beyond the sites (to the north or higher in elevation). The sites are underlain by continuous permafrost and tend to be quite wet. The location is characterized by cold temperatures (annual average = -8.1 °C) and modest precipitation (485.4 mm yr^-1^), as measured from a nearby Snow Telemetry (SNOTEL) site (https://wcc.sc.egov.usda.gov/nwcc/site?sitenum=957).

The six sites were established along the Dietrich River flood plain in 2016 (Eitel *et al*., 2019). At each site, a circular plot with 10 m radius (314 m^2^ area) was established, and six white spruce trees from two different size classes were tagged. The size classes were chosen according to Körner (2012b) to differentiate “trees” (≥ 10 cm DBH) from “saplings” (<10 cm DBH).

In June 2018, leaves were sampled from two locations for each of the 36 target trees, made up of 18 saplings and 18 trees, for gas exchange analysis. The terminal portions of several branches were cut with sharp pruners (bottom of the canopy) or a pole clipper (top of the canopy), and the removed portion of the stem was immediately wrapped with wet paper towels, sealed in a plastic bag with ample air and placed in a cooler where they were kept in the dark until arriving in the lab. The location of the lower samples was approximately 1.37 m above the ground, and were collected from the south side of the canopy. The location of upper samples was approximately 1 m from the terminal apex of each tree, and were collected from the south side of the canopy. Once returned to the lab, the stems were recut underwater and placed in a beaker containing enough water to keep the cut end submerged until analyzed, typically within 8, but no more than 24 hours.

### Respiration Temperature Response Curves

To assess average respiration rates at a common temperature (25°C, *R*_25_) and to quantify the response of respiration to temperature, CO_2_ exchange rates were measured. The individual leaves (needles) were carefully removed from the stems to avoid the large contribution the stem would otherwise have made to the CO_2_ flux thereby confounding the results (e.g. (Heskel *et al*., 2014)). These needles were weighed to determine the initial fresh mass (g) and then placed in a fine nylon mesh bag, allowing for easy air flow through the bag while keeping the leaves from entering the optical bench of the infrared gas analyzer or becoming lost in the leaf cuvette. The mesh bag containing the leaves was placed inside a custom-made cuvette milled from a solid block of aluminum with a plexiglass lid sealed with a Viton gasket (Patterson *et al*., 2018; Li *et al*., 2019; Schmiege *et al*. 2021)). The cuvette contained a mixing fan and two fine wire thermocouples to measure leaf and air temperatures. Temperature was thermoelectrically controlled from a laptop computer (using a CP-121 Thermoelectric Peltier Cooling Unit, TE Technology, Traverse City, MI USA). The custom cuvette was interfaced with a portable photosynthesis system (Li-6800XT, LiCor Lincoln, Nebraska USA).

Before each measurement, the system was both equilibrated to 5°C and zeroed. The mesh bag holding the leaves was then sealed inside the cuvette and the system was again equilibrated to 5°C. Once stability was reached, the response curve was measured as described in previous studies (O’Sullivan *et al*., 2013; Heskel *et al*., 2016; Schmiege *et al*. 2021). Conditions during the measurements included a flow rate of 500 ml min^-1^ through the cuvette, and a reference CO_2_ concentration of 400 ppm. The incoming air was dried using the Li-6400XT desiccant column and transpiration was allowed to humidify the air as done previously (*see references above*). During the measurement the cuvette temperature was ramped continuously from 5 to 65°C at a constant rate of 1°C min^-1^. All gas exchange variables and calculated rates were recorded every 20 seconds by the Li-6400XT.

### Leaf Traits

Upon the completion of the temperature response curve, the leaves were removed from the cuvette and photographed with a known scale. ImageJ software (Schneider *et al*., 2012) was used to determine the projected area of these needles. The leaves were transferred to coin envelopes and dried at 65°C for a minimum of 48 hours. Dried leaves were again weighed to determine leaf dry mass (g). Specific leaf area (SLA cm^2^ g^-1^) was calculated and used to interconvert between area- and mass-based respiratory fluxes. Leaf water content (%) and leaf dry matter content (LDMC) were calculated from the fresh and dry masses. Leaf nitrogen was estimated using the %N measured from these same trees, sampled at the same canopy locations in 2017 (Schmiege *et al. unpublished data*).

### Data Analyses

The respiration temperature response curves were analyzed as in Heskel *et al*. (2016) by fitting a second-order polynomial model to the log transformed respiration rates between 10 and 45 °C:

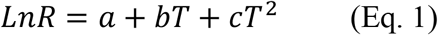

Where *a* represents the basal respiration rate (y-intercept), and *b* and *c* describe the slope and curvature of the response (Heskel *et al*., 2016). From these modeled curves we also derived the respiration rate at a common temperature of 25°C (*R*_25_). Using the entire data set between 5 and 65°C the temperature of the maximum respiration rate was extracted (*T*_max_).

Before statistical analysis, all traits were transformed as necessary to fulfill statistical assumptions of normality. Statistical differences in *P. glauca* traits between large trees and small saplings including DBH and tree height were assessed using an independent sample t-test. To test our hypothesis regarding the main effects of canopy position or tree size on all respiratory parameters (including the polynomial model parameters *a, b*, and *c*, area- and mass-based *R*_25_, and *T*_max_) and leaf traits (including SLA, LDMC and %N), we used a linear mixed effects model allowing us to incorporate each unique tree as a random effect (using the lme4 and the lmerTest packages; Bates *et al*., 2015; Kuznetsova *et al*., 2017, respectively). Finally, model coefficients from our study were compared to those from Heskel *et al*. (2016) for the Tundra and Boreal biomes, as well as the needle-leaved evergreen (NLEv) plant functional type using the information contained within their Table 1, with an independent sample t-test. For all analyses, statistical significance was assessed using a p-value of 0.05. Data analysis was done in either R version 3.6.3 (R Core Team, 2020) or Microsoft Excel®.

**Table 1.** Tree and leaf characteristics of *Picea glauca* trees (≥ 10cm DBH – dark green) and saplings (5-10cm DBH – light green) growing in the Forest Tundra Ecotone in Alaska. DBH = diameter at breast height (1.37m), SLA = specific leaf area, LDMC = leaf dry matter content (g dry mass g^-1^ fresh mash, and N = leaf nitrogen %. ^*^ Leaf nitrogen data collected from the same trees in 2017, by Schmiege. n = 36 for both trees and saplings, 72 for all trees.

|  | *DBH* | *Height* | *SLA* | *LDMC* | *N* * |
| --- | --- | --- | --- | --- | --- |
|  | *cm* | *m* | *cm<sup>2</sup> g<sup>-1</sup>* |  | *%* |
| Saplings | 6.58 ± 0.25 **a** | 4.18 ± 0.18 **a** | 21.99 ± 0.35 **a** | 0.54 ± 0.004 **a** | 0.93 ± 0.02 **a** |
| Trees | 16.60 ± 0.68 **b** | 9.36 ± 0.41 **b** | 21.84 ± 0.35 **a** | 0.54 ± 0.004 **a** | 0.99 ± 0.02 **a** |
| All | 11.52 ± 0.69 | 6.73 ± 0.38 | 21.90 ± 0.22 | 0.54 ± 0.002 | 0.95 ± 0.66 |

## RESULTS

### Tree and leaf traits

The average tree in this study was 11.52 ± 6.73 cm in DBH (1.37 m from the ground, Table 1). When divided into two size classes, the saplings (5-10 cm DBH) were on average 60% (∼ 10 cm) smaller than the trees (≥10 cm DBH) (Table 1). In addition, trees were 55% taller than the sapling class. None of the measured leaf traits differed by size class (trees vs. saplings, Table 1).

### Respiration temperature response curves

All leaves showed a similar response to temperature during the high-resolution measurements, increasing exponentially between approximately 5 and 45 °C, then slowing briefly before increasing with rising leaf temperatures to a maximum rate of respiration at *T*_max_ (Figure 1a). To model the ecologically relevant response, the data were limited to between 10 and 45°C and the log normal response was fit with the global polynomial model of (Heskel *et al*., 2016) (Figure 1b, equation 1 above). This model fit all curves remarkably well with r^2^ ≥ 0.99 in all cases. Overall, the three model (see Eq. 1) coefficients averaged -0.931 ± 0.069, 0.083 ± 0.002 and -0.00023 ± 5.5 × 10^−5^ (*a, b* and *c* respectively, mean ± SEM). There was no statistically significant effect of canopy position on any of the model fits, model coefficients or derived variables (data not shown). However, saplings did differ significantly from trees (Table 2, Figure 2). The intercept (coefficient *a*) of the lnR/T response of saplings was 38% lower than basal respiration for the trees. Neither the slope nor the curvature (coefficients *b* & *c*) of the lnR/T relationships differed between trees and saplings (Table 2, Figure 2). The response of white spruce respiration to varying temperature differs from the average response for species in the needle-leaved evergreen plant functional type and both the Tundra and Boreal biomes in the global compilation of Heskel *et al*. (2016) (Figure 3). In all cases the *a* coefficient of both our trees and saplings differed significantly from the *a* coefficient of the Tundra or Boreal Biomes, and the NLEv PFT of Heskel *et al*. (2016) (*p*<0.5). While the slope and curvature (*b* & *c* coefficients) were statistically similar for the comparisons of either the FTE trees or saplings with the Boreal Biome, both FTE groups differed from the Tundra biome and NLEv PFT.

**Figure 1.**
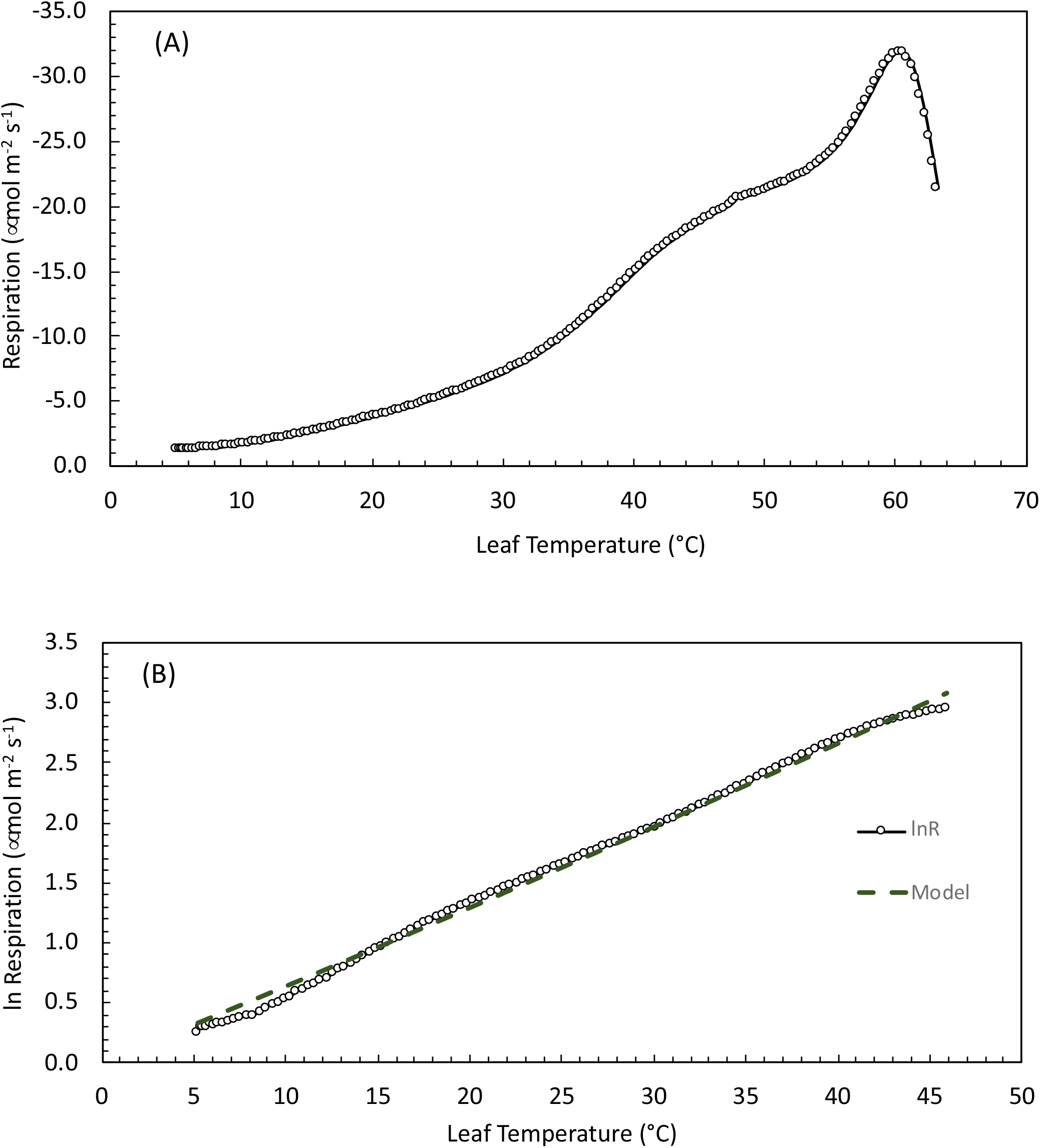
Leaf respiration as a function of temperature in *Picea glauca* measured at the Forest Tundra Ecotone in Alaska. Panel A (top) is an example of the high-resolution temperature response curves measured from 5 to 65°C. Air temperature was heated at a rate of 1°C min^-1^while the rate of CO_2_ release and other gas-exchange parameters were recorded every 20 seconds. Panel B (bottom) is the log of measured respiration rate and model fit (*lnR = a + bT + cT*^*2*^ – see text for full description) between 10 and 45 °C (the ecologically relevant temperature range) for the same sample.

**Table 2.**
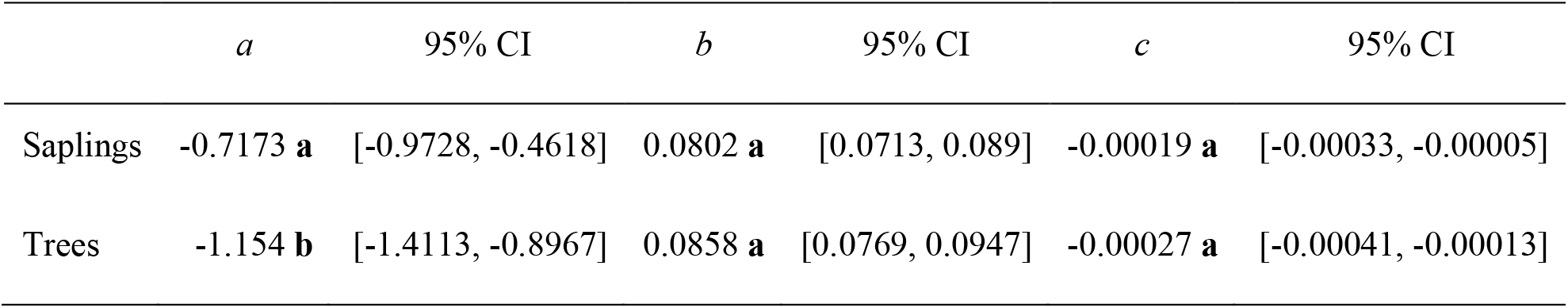
Model parameters fit to the measured high-resolution leaf respiration temperature response curves collected from *Picea glauca* measured at the Forest Tundra Ecotone in Alaska. Results are separated by trees (≥ 10cm DBH) and saplings (5-10cm DBH) trees and saplings. Mean response (n=36) and 95% confidence interval for each size class. Differing letters following the means denote statistically significant differences.

|  | *a* | 95% CI | *b* | 95% CI | *c* | 95% CI |
| --- | --- | --- | --- | --- | --- | --- |
| Saplings | -0.7173 **a** | [-0.9728, -0.4618] | 0.0802 **a** | [0.0713, 0.089] | -0.00019 **a** | [-0.00033, -0.00005] |
| Trees | -1.154 **b** | [-1.4113, -0.8967] | 0.0858 **a** | [0.0769, 0.0947] | -0.00027 **a** | [-0.00041, -0.00013] |

**Figure 2.**
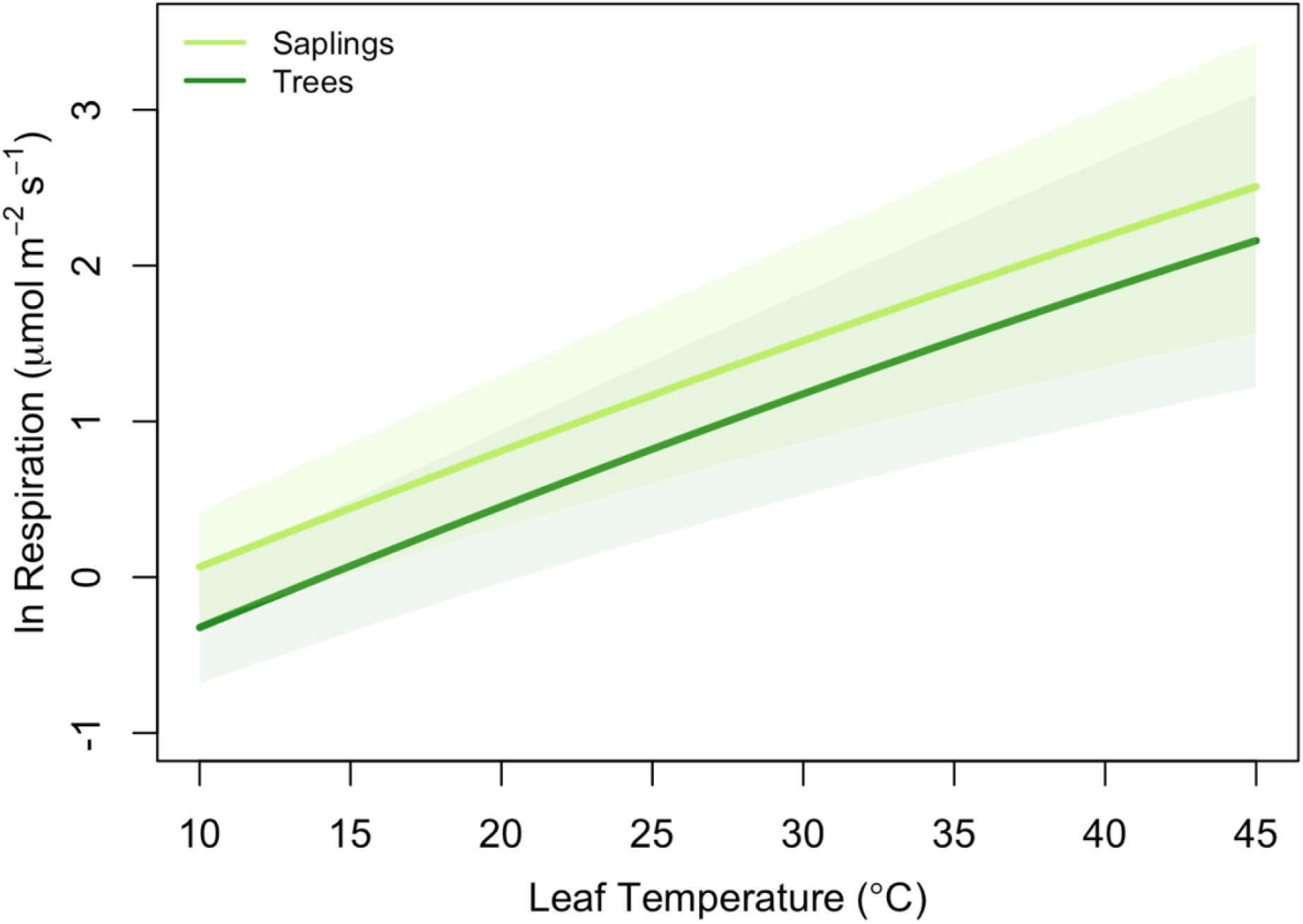
Average model results (log of leaf respiration) for trees (≥ 10cm DBH – dark green) and saplings (5-10cm DBH – light green) of *Picea glauca* growing in the Forest Tundra Ecotone in Alaska. Line presents the mean response (n=36) and shaded area = 95% confidence interval for size class. Model parameters are presented in Table 2.

**Figure 3.**
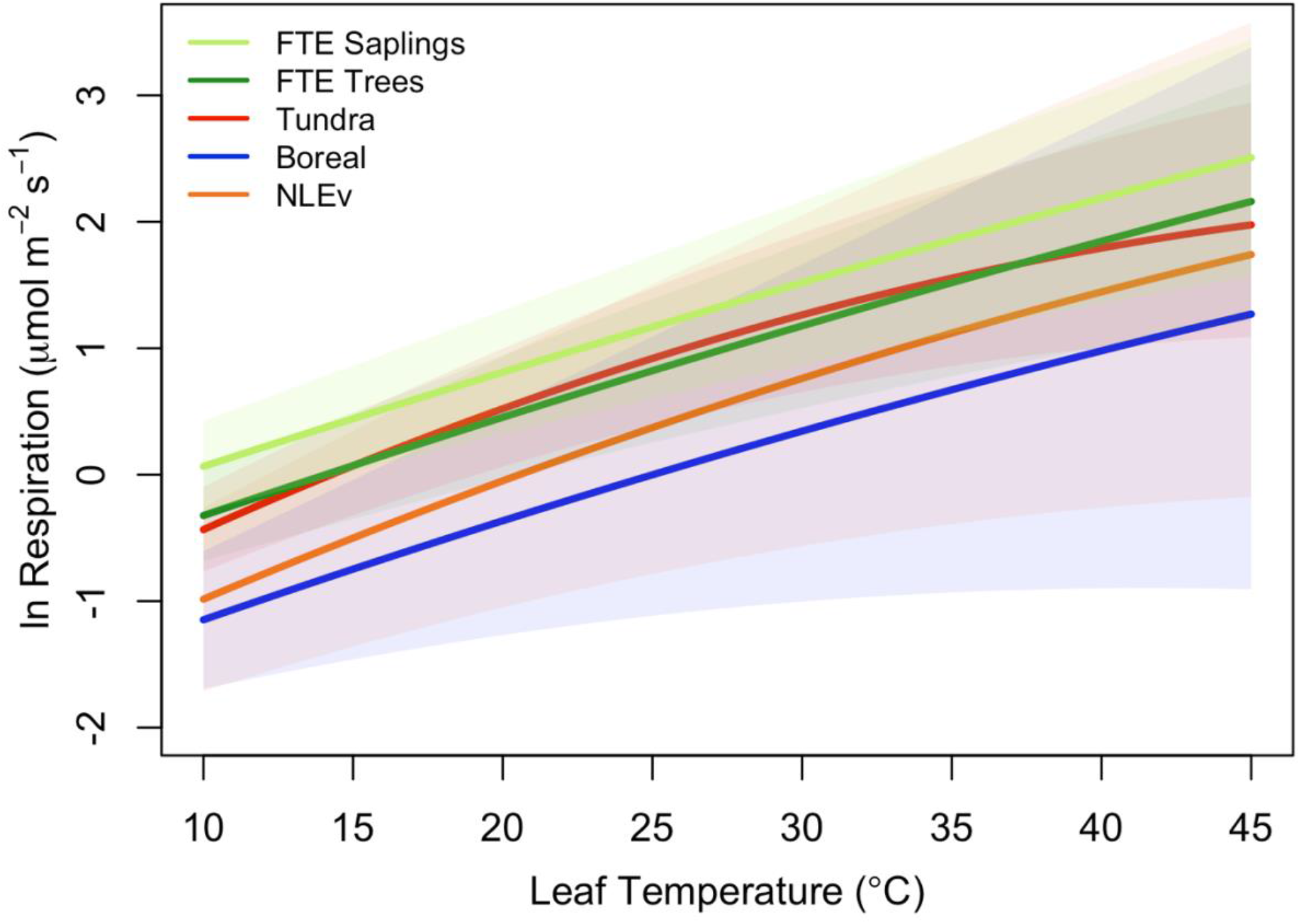
Comparison of Polynomial models fits from Heskel *et al* (2016, 2019) with the average fits measured on two size classes of *Picea glauca* at the FTE in Northern Alaska. Response plotted are for the tundra biome (red line), boreal biome (blue line), needle-leaved evergreen plant functional type (orange line), FTE trees (>10cm; dark green line), and FTE saplings (5-10 cm DBH; light green line).

### Respiration at a common temperature & *T*_*max*_

Across all samples, the average rate of respiration at a common temperature of 25°C was 2.92 ± 0.135 μmol CO_2_ m^-2^ leaf area s^-1^. This rate did not differ by canopy position but did vary between saplings and trees. Saplings had a significantly higher area-based *R*_25_ than larger trees (3.43 ± 0.19 μmol m^-2^ s^-1^ vs. 2.41 ± 0.14 μmol m^-2^ s^-1^, *p*<0.5) (Figure 4a). The 43% lower area-based rate observed in the tree *R*_25_ was similar when converted to a mass basis (0.00738 ± 0.00036 μmol g^-1^ s^-1^ vs. 0.00525 ± 0.00032 μmol g^-1^ s^-1^ in saplings vs. trees respectively, Figure 4b). Saplings respire at leaf temperatures that were nearly 2°C higher than the *T*_max_ for large trees (57.6°C, Figure 5). None of the respiratory parameters (*R*_25_, *a* or *T*_max_) were significantly related to leaf N (mg N m^-2^) (F significance = 0.13, 0.18 and 0.72 respectively, data not shown).

**Figure 4.**
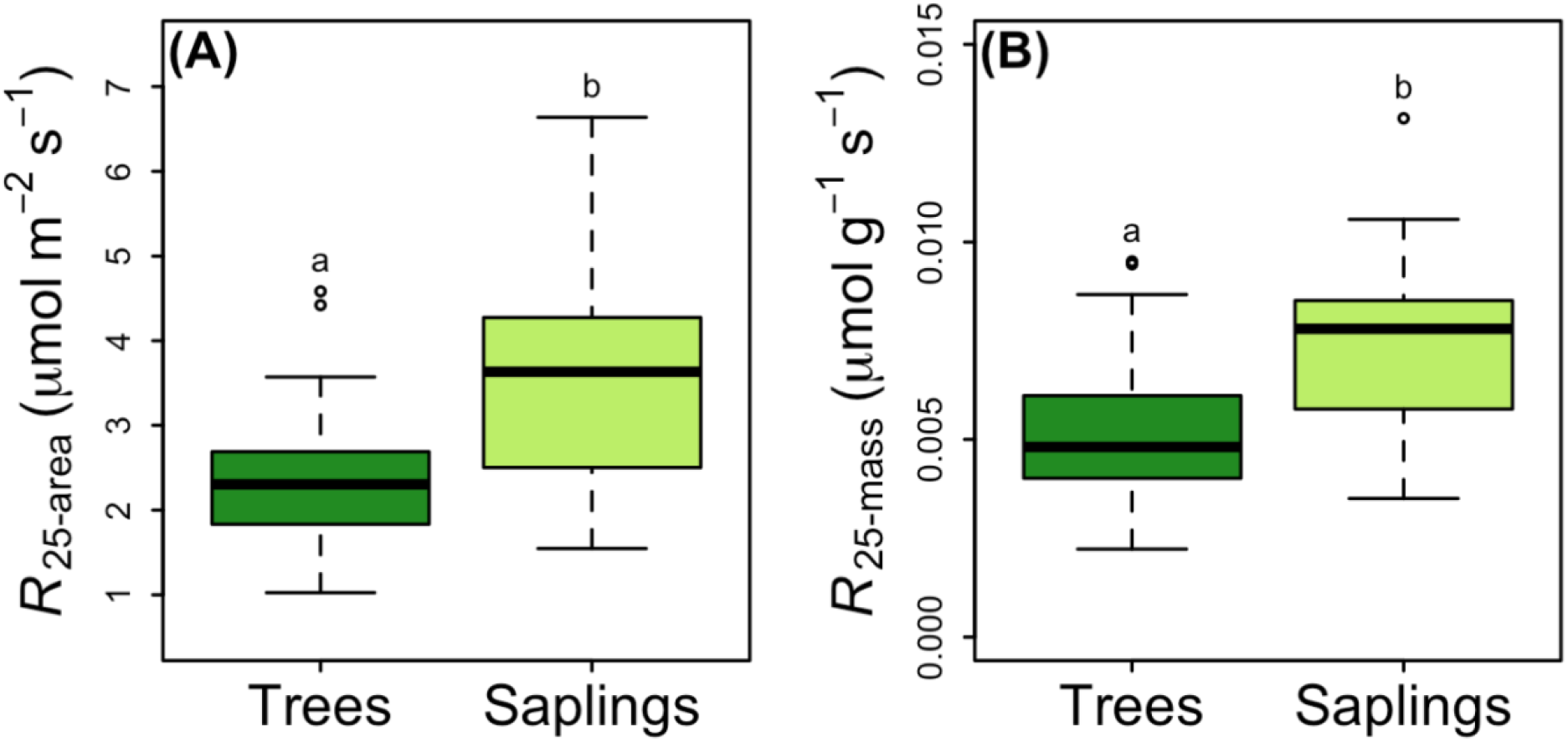
*Picea glauca* leaf respiration normalized to 25°C, as a function of tree size at the Forest Tundra Ecotone in Alaska. Respiration is expressed as a function of leaf area (panel A, left) and leaf mass (panel B, right). Boxplots represent the median and the first and third quartiles. Whiskers delimit the range for each group, with outliers falling outside 1.5 times the interquartile range marked by points. Significant differences between groups are marked by different letters (p < 0.05). Dark green boxes = trees ≥10 cm DBH, light green boxes = saplings 5 - 10 cm DBH. n = 36.

**Figure 5.**
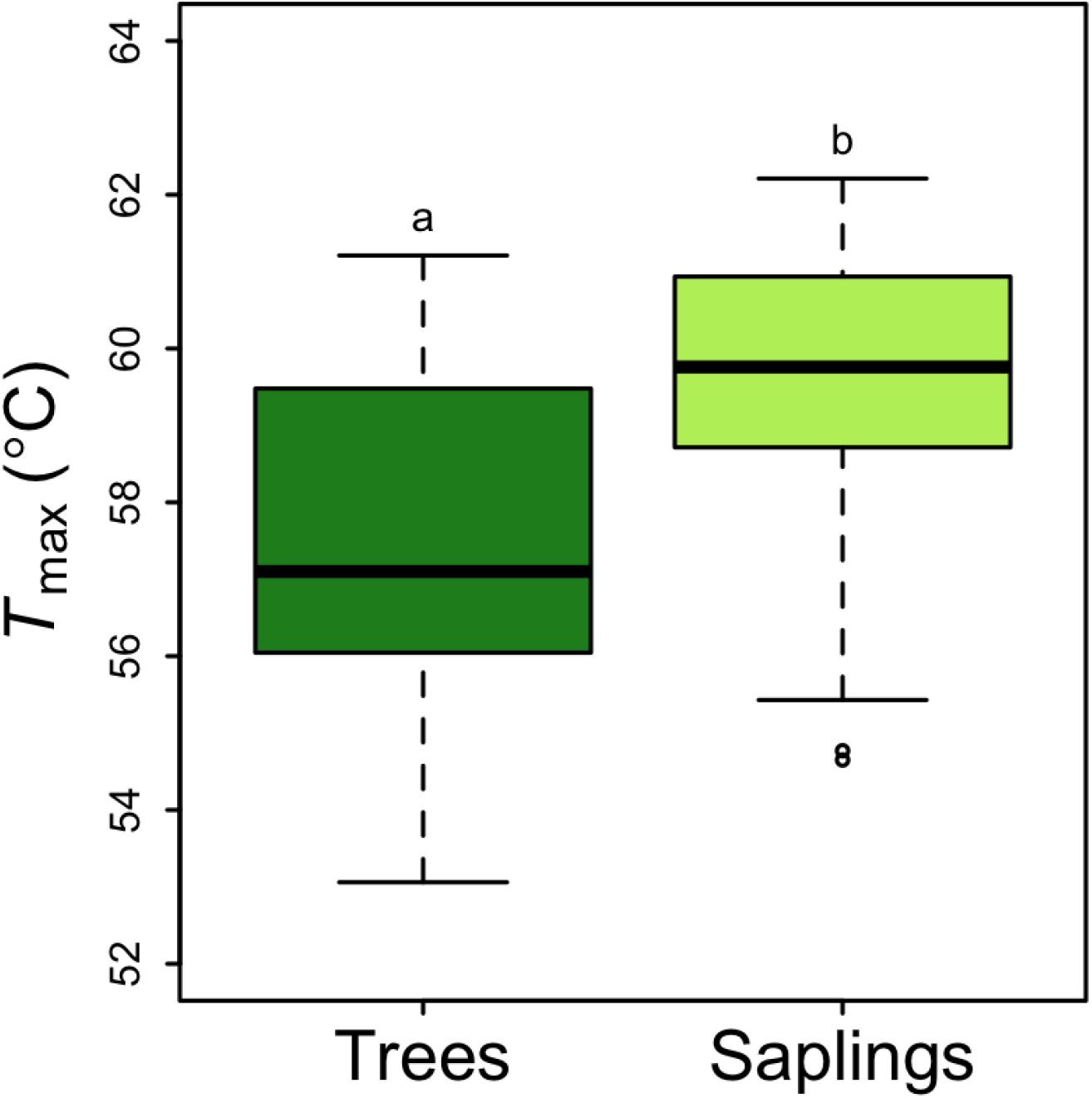
*Picea glauca T*_max_ (the leaf temperature corresponding to the maximum rate of respiration was reached), as a function of tree size at the Forest Tundra Ecotone in Alaska. Boxplot represent the median and the first and third quartiles. Whiskers delimit the range for each group, with outliers falling outside 1.5 times the interquartile range marked by points. Significant differences between groups are marked by different letters (p < 0.05). Dark green box = trees ≥10 cm DBH, light green box = saplings 5 - 10 cm DBH. n = 36.

## DISCUSSION

We show that saplings have significantly higher rates of respiration than trees. This result supports our first hypothesis (H_01_/H_A1_) and is highly important as it ultimately may help explain the mechanistic underpinnings of the structure and function of the Forest Tundra Ecotone (FTE). Any discussion of the FTE, the role of climate in determining its location, or the effect of climate change on its rate of migration, must begin with a definition of ‘tree’. Here we adopt the terminology of Körner (2012a & b), who presents the following life stages: *Germlings*, in the year of germination; *Seedlings*, commonly < 15 cm tall; *Saplings*, with stem of less than 10 cm DBH, and finally; *Trees*, with DBH greater than 10 cm. Since treeline, and hence the FTE, is defined by the lack of trees, the implication is that germlings, seedlings and saplings growing at this transitional boundary between tundra and forest (the *kampfzone*), do not represent the permanent establishment of the boreal forest biome, as they are unlikely to survive. The question pondered by tens, perhaps hundreds of scientists, is – why are they unlikely to survive? Are there environmental and/or biological conditions that limit the transition to the tree lifeform? While dozens of possible explanations have been studied, and many variables deemed to contribute (locally), cold temperatures have always been the most likely constraint on the position of *global* treeline (both alpine and northern) (Korner, 1998; Korner & Paulsen, 2004; Körner, 2012b). Our data provide mechanistic, physiological evidence that saplings incur higher respiratory costs without displaying thermal acclimation (compared to trees). These higher costs may result in carbon balance limitations that ultimately limit sapling survival.

White spruce is known as an extremely hardy and robust species that can persist for hundreds of years while growing very slowly (Sutton, 1969; Nienstaedt 1990). Slow growth can be part of a relatively common life history strategy that includes low photosynthetic rates and large investments in long lived leaves, and thus results in long returns on carbon investments (Poorter & Pothmann, 1992; Poorter *et al*., 2014; Reich, 2014). Adding high basal metabolic costs (high respiration rates) to this strategy could help explain the difficulty FTE saplings have in surviving the harsh conditions and short growing season. Previous work on white spruce (Man & Lieffers, 1997; McNown & Sullivan, 2013; Man & Lieffers 1997; Stinziano & Way, 2017; Benomar *et al*., 2018; Prud’homme *et al*., 2018) demonstrate that it has, at best, modest maximum photosynthetic rates (typically less than 10 μmol m^-2^ s^-1^) and that these rates may be realized only rarely when the ambient environmental conditions are most favorable. Furthermore, McNown & Sullivan (2013) also present data from treeline in Northern Alaska, demonstrating that the net photosynthesis of white spruce can average less than 4 μmol m^-2^ s^-1^ while leaf respiration can exceed 2.5 μmol m^-2^ s^-1^. This leaves an exceptionally thin carbon margin to support growth and metabolic function (McNown & Sullivan, 2013, Figure 3). Unpublished data from our site (Schmiege *et al*.), find higher rates of net photosynthesis (collected in 2017, the year before the current respiration measurements) than those of McNown and Sullivan (2013), yet do not preclude the limiting carbon balance mechanism. In fact, the *R*_25_ rates for saplings presented here average 3.4 μmol m^-2^ s^-1^,which are significantly higher than both those reported by McNown & Sullivan (2013), and the global average for needle-leaved trees (∼1.25 μmol m^-2^ s^-1^, GlobResp database, Atkin *et al*., (2015)) demonstrating the potential for exceptionally high respiratory costs for saplings growing and attempting to establish themselves as trees in the FTE. Interestingly, individual plants with DBH <10cm (conventionally called ‘saplings’) are routinely >100 years old at the FTE, further complicating assumptions about tree demography and implications of relatively rapid warming-induced respiratory losses for this critical size class.

Respiration at a common temperature (*R*_25_) is a useful tool for comparing across studies, across ecological settings and across biomes (*e*.*g*., Atkin *et al*. 2015), but alone, is not a sufficient indicator of *in situ* respiratory activity. High-precision temperature response curves, like those presented here, can provide significantly more information and insight into how trees respond to changing local environmental conditions (O’Sullivan *et al*., 2013). Furthermore, coupling these measurements to models, such as the global polynomial model of Heskel *et al*. (2016) provides the tools for analyzing, comparing and scaling the results (Huntingford *et al*., 2017). We find the intercept of the relationship between leaf temperature and the log of respiration (coefficient *a*), differs significantly between trees and saplings. This finding extends the *R*_25_ result and again suggests a higher respiratory cost for saplings, even at cool (< 10 ° C) temperatures. Not only is the cost higher for FTE saplings, but overall, the costs are high compared to other needle-leaved evergreen, boreal or tundra plants of all sizes. For example, Heskel *et al*’s (2016) global survey reports the average *a* coefficient to be -2.05 across all needle-leaved evergreen trees, -2.00 in the boreal biome and -1.6 in the Tundra biome, while we report -1.2 for trees and -0.7 saplings at the FTE. In contrast to these differences in *a*, there were no differences in either the slope (*b*) or curvature (*c*) of the modeled temperature responses between the two size classes at the FTE, which is in contrast to our hypothesis. The lack of change in these coefficients suggests that as temperatures change (over the course of the day, the season or perhaps even over the years), respiratory rates will not be moderated and will remain high across all temperatures. This pattern is functionally similar to the theoretical “Type II acclimation” (an upwards shift in low temperature respiration, or coefficient *a*, without changes in the overall shape (*b* & *c*), (Atkin & Tjoelker, 2003)), which has been attributed to limited respiratory enzyme capacity. The shape of the FTE R/T response (as defined by the *b* & *c* coefficients of the model) also deviates significantly from the average responses of both the Tundra biome and NLEv PFT, it is similar to that of the Boreal biome (collected with similar techniques, Heskel *et al*., (2016)). The high respiratory costs of saplings at the FTE are thus unique (Heskel *et al*., 2016) and may be rooted in the limited capacity of their respiratory enzymes. Any number of environmental and biological conditions may contribute, but clearly across ecologically relevant temperatures, the FTE saplings have higher respiratory carbon losses that must be overcome to survive.

What drives higher respiratory rates in saplings compared to trees at the FTE, and in white spruce at the FTE compared to other species of similar functional types, or growing in neighbouring biomes more generally? We have only limited ability to answer this question, but find no evidence of differences in nitrogen nutrition (large trees had similar leaf nitrogen concentrations to saplings and we found no relationship between leaf N and *R*_25_ or *a*), water availability (similar % leaf water content) or structural investment (similar LDMC, SLA and consistent differences in area and massed based respiration rates). Instead, we suggest that faster relative growth rates of the saplings at this site (Jensen *et al*., unpublished data), and presumably higher maintenance costs, both contribute. Additional studies designed to separate growth and maintenance respiration (Amthor, 1984) would provide insight, as would detailed quantification of the microenvironment as a function of tree size (Scott *et al*., 1987; Moser *et al*., 2010; Smith *et al*., 2009; Körner, 2012b; Maguire, A. J. *et al*., 2019), with particular focus on the interactions among temperature, windspeed and vapor pressure deficit (*e*.*g*. Holmgren *et al*., 1996; Squeo *et al*., 1991; Holtmeier, 2009; Maguire *et al*., 2019).

An additional piece of information available in the high-resolution temperature response curves is *T*_max,_ the temperature at which the highest respiration rate was achieved. This temperature is related to leaf death, and at temperatures exceeding *T*_max_, respiration drops precipitously and irreversibly (O’Sullivan *et al*., 2013). Our estimates of *T*_max_ are quite high but within the upper bounds of the global survey of O’Sullivan *et al*., (2017). However, our results stand out from the latitudinal patterns found by O’Sullivan *et al*. (2017) that clearly show a decreasing *T*_max_ at higher absolute latitudes and thus would predict much lower *T*_max_ values at the FTE. In fact, the *T*_max_ we report here for white spruce within the FTE is much closer to the lowest absolute latitude samples and highest maximum temperature sites from the cross-biome survey (O’Sullivan *et al*., 2017). This earlier work concluded that plants at mid-latitude sites were operating closest to, and sometimes outside of, the respiratory thermal safety margin (e.g. the difference between *T*_max_ and the mean maximum daily temperature over the warmest consecutive 3-day period) and were most at risk for mortality during heat waves (O’Sullivan *et al*., 2017). By contrast, plants growing at either high or low latitudes were found to have the highest margin of safety and were unlikely to be damaged by heat wave events. In comparison, we find that the thermal safety margin at the FTE is curiously high, suggesting that white spruce growing at treeline are at very low risk for thermal damage, even during extreme heat waves with temperatures as unrealistically high as 59°C. The reason for this high temperature tolerance is unknown, but perhaps related to leaf morphology, tree age (having experienced a greater range of temperatures), or the magnitude of the respiratory “burst” (Hueve *et al*., 2011; Huve *et al*., 2012; O’Sullivan *et al*., 2017). The respiratory burst is thought to emanate from heat-induced changes in membrane properties and the rate of ATP synthesis and/or demand (Berry & Bjorkman, 1980; Seemann *et al*., 1984; Hazel, 1995; Sung *et al*., 2003; Zhu *et al*., 2018). While heat wave events may not challenge white spruce directly, indirectly the effects of rising temperature on leaf and tree carbon balance could still hamper sapling survival and provide the mechanism linking treeline locations to temperature as suggested by Körner (2012b).

Warming in the Arctic is dramatic and occurring much faster than at lower latitudes (Huang *et al*., 2017). Furthermore, the rate of warming will be exacerbated by continued greenhouse gas emissions. Predictions for the interior of Alaska are for an additional temperature increase of up to 6°C by the end of the century (Markon *et al*., 2012). As thermal acclimation of respiration has been shown to be limited in white spruce (Benomar *et al*., 2018), respiration rates within the FTE could increase significantly under these scenarios. Currently June, July and August temperatures average approximately 15°C (Harris *et al*., 2020; Zepner *et al*., 2021). If these average temperatures were to increase to 21°C, we estimate that leaf respiration would increase by 55%, from 1.07 to 1.70 μmol m^- 2^ s^-1^ in trees and 1.56 to 2.42 μmol m^-2^ s^-1^ in saplings. Despite the potential for photosynthesis to acclimate to higher temperatures, Stinziano & Way (2017) found that with warming, white spruce allocates additional carbon to respiration rather than to growth. We conclude that this carbon balance challenge may be particularly acute at the FTE. Further studies of the physiological mechanisms involved, such as photosynthesis, respiration and growth, are urgently needed.

## Conclusions

In his influential book on treeline, Körner (2012b) describes the sapling stage as being the most critical for woody plant survival at or near treeline.

> “After seedling establishment, the most critical life stage is the emergence from the aerodynamic boundary provided by alpine heathland or by micro-topography to the open convective conditions in the free atmosphere. This transition is a continuous ‘fight’ that led to the German term ‘kampfzone’, describing the uppermost belt of the ecotone, where trees quite often lose the fight and remain crippled, forming shrub-like structures termed ‘krummholz’. Perhaps, this is the central question of treeline formation, what causes tree species to remain confined to the shrub layer rather than to grow into upright trees. (Page 122)”

Here we show that saplings struggling to grow beyond the aerodynamic boundary layer have higher rates of respiration and thus incur a carbon cost beyond what is present in established trees. The higher respiration rate is not accompanied by a change in the shape of the temperature response curve and thus the additional cost, above that of the trees, will be incurred at all temperatures. These findings partially support our second hypothesis (H_02_), that tree size does not influence leaf respiration. However, we found no support for our first hypothesis on the effect of canopy position. Respiratory characteristics of leaves at the top of the canopy were no different from those of leaves at the bottom of the canopy. We have demonstrated that compared to previous measurements from the Tundra or Boreal biomes, as well as the global average of the needle-leaved evergreen plant functional type, the needles of white spruce growing at the Alaskan FTE are respiring at what are likely the species extremes as demonstrated by their very high respiratory rates at a common temperature, their high measured *T*_max_, and the lack of acclimation in the shape of temperature response of respiration. Our findings suggest the capacity of respiratory enzymes could be limiting the rate of respiration of this species at treeline. Without thermal acclimation, respiratory carbon release could be as much as 57% higher by the end of the century and potentially preclude the predicted (but not often observed) advance of the FTE. A more robust examination of the physiological processes contributing to leaf and tree carbon balance at the FTE is urgently needed.

## AUTHOR CONTRIBUTIONS

K.L.G., S.C.S., N.B., L.A.V. and J.U.H.E. designed the research. They were assisted in data collection by S.G.B.. S.C.S & K.L.G. analyzed the data. K.L.G wrote the first draft of the manuscript. All authors contributed to the revisions, editing and submission of the final manuscript.

## ACKNOWLEDGMENTS

We thank Sarah Sackett from the NASA ABoVE support team in Fairbanks, AK, for logistical support during Alaska field campaigns. We thank Zoe M. Griffin for lab assistance.

## DATA AVAILABILITY STATEMENT

The datasets, generated for this study can be found in the Oak Ridge National Laboratory Distributed Active Archive Center at: *XXXXXXXX (note we are currently in the process of archiving the data)*

